# Forecasting the effects of water regulation on the population viability of a threatened amphibian

**DOI:** 10.1101/2021.04.20.440713

**Authors:** Rupert Mathwin, Skye Wassens, Matthew S. Gibbs, Jeanne Young, Qifeng Ye, Frédérik Saltré, Corey J.A. Bradshaw

## Abstract

The regulation of river systems alters hydrodynamics and often reduces lateral connectivity between river channels and floodplains. For taxa such as frogs that rely on floodplain wetlands to complete their lifecycle, decreasing inundation frequency can reduce recruitment and increase the probability of local extinction. We virtually reconstructed the inundation patterns of wetlands under natural and regulated flow conditions and built stochastic population models to quantify the probability of local extinction under different inundation scenarios. Specifically, we explored the interplay of habitat size, inundation frequency, and successive dry years on the local extinction probability of the threatened southern bell frog *Litoria raniformis* in the Murray River floodplains of South Australia. We hypothesised that the changes to wetland inundation resulting from river regulation are a principal driver of *L. raniformis* declines in this semi-arid system.

Regulation has reduced the inundation frequency of essential habitats below critical thresholds for the persistence of many fresh water-dependent species. Successive dry years raise the probability of local extinction, and these effects are strongest in smaller wetlands. Larger wetlands and those with more frequent average inundation are less susceptible to these effects.

Elucidating these trends informs the prioritisation of treatment sites and the frequency of conservation interventions. Environmental water provision (through pumping or the operation of flow-regulating structures) is a promising tool to reduce the probability of breeding failure and local extinction. Our modelling approach can be used to prioritise the delivery of environmental water to *L. raniformis* and potentially many other frog species.

## Introduction

The global decline of amphibians (Blaustein and Wake 1990; Stuart et al. 2004) is stark, with the most recent figures showing 41% of assessed species are threatened with extinction (IUCN 2020). Human modifications of wetland networks on which many amphibian populations depend via river regulation can reduce lateral connectivity and the ecological function of wetland habitats (Castello and Macedo 2016). Being essential breeding and nursery areas for many amphibian species, such regulation can change amphibian community structure and drive local extinctions (Wassens and Maher 2011). Furthermore, amphibian extinctions and population declines are projected to increase through the 21^st^ Century as the interactive perturbations increase in intensity (Hof et al. 2011); for example, the negative consequences of river regulation on amphibians are predicted to increase as climate change progressively overtakes land use (Narins and Meenderink 2014) as the main driver of species richness patterns and extinctions (Newbold 2018). Climate change-driven aridification reduces the availability of breeding habitats, resulting in lower species richness (McMenamin et al. 2008). The combination of aridification and increasing water consumption is therefore tlikely to reduce the availability of freshwater habitats further in some regions (Miller et al. 2018).

Manipulating water resources can help alleviate the effects of declining water availability and support amphibian recruitment (Shoo et al. 2011; Smith et al. 2019; Mathwin et al. 2021). Techniques for manipulating water to benefit amphibians vary in approach and success (Mathwin et al. 2021), but the best-supported intervention is the provision of water to breeding habitats to match the larval requirements of the target species (‘environmental water provision’). This is because enhancing survival through breeding and early life stages can stabilise populations (Griffiths and Pavajeau 2008). The targeted delivery of environmental water is a common practice today (Kennen et al. 2018) and could become necessary to conserve some species (Greenwood et al. 2016), especially those with limited phenotypic plasticity or those near the edge of their ecological niche (Grant et al. 2020).

Australia’s Murray-Darling Basin provides a model system to examine this process. The catchment is heavily regulated and each year up to 61% of total flow is extracted for consumptive use (CSIRO 2008). This has reduced the number and function of wetlands (Gell and Reid 2014), resulting in the decline and fragmentation of water-dependent taxa (including the southern bell frog, *Litoria raniformis*) (Clemann and Gillespie 2012). In response to systemic environmental degradation, federal legislation mandates interventions aimed at restoring ecological function (Docker and Robinson 2014), including environmental water provision for species recovery.

In this paper we explore hydro-ecological thresholds and generate guidelines for environmental water provision to benefit amphibians by constructing stochastic, hydro-ecological population models. We hypothesise that the reduced frequency of wetland inundation resulting from river regulation is driving local extinction events of *L. raniformis* in this system. We hypothesise that larger wetlands and wetlands with greater average inundation are less susceptible to local extinction during successive dry years. We posit that the probability of local extinction can be reduced by environmental water provision and that our modelling approach can be used to prioritise the location and frequency of environmental water provision to at-risk amphibian populations.

## Materials and Methods

### Study area

The Murray-Darling Basin contains Australia’s longest river system and is heavily regulated to provide water for domestic and agricultural use. Regulating structures include a series of 14 main-channel weirs (‘locks’) that dissect the river. Our focus is 70 km of the Murray River channel between Lock 3 (34° 11′ 16.95″ S, 140° 21′ 29.65″ E) and Lock 2 (34° 4′ 39.31″ S, 139° 55′ 52.81″ E) and the associated wetlands and floodplains along this reach (**Figure 1**). This region receives an annual rainfall of 160–240 mm (semi-arid/arid), which is insufficient to fill off-channel wetlands most years. As such, all naturally occurring *L. raniformis* breeding in this reach results from elevated river level or the provision of environmental water. Water is typically delivered using portable pumps that exclude most fish and aquatic invertebrate predators.

**Figure 1.**
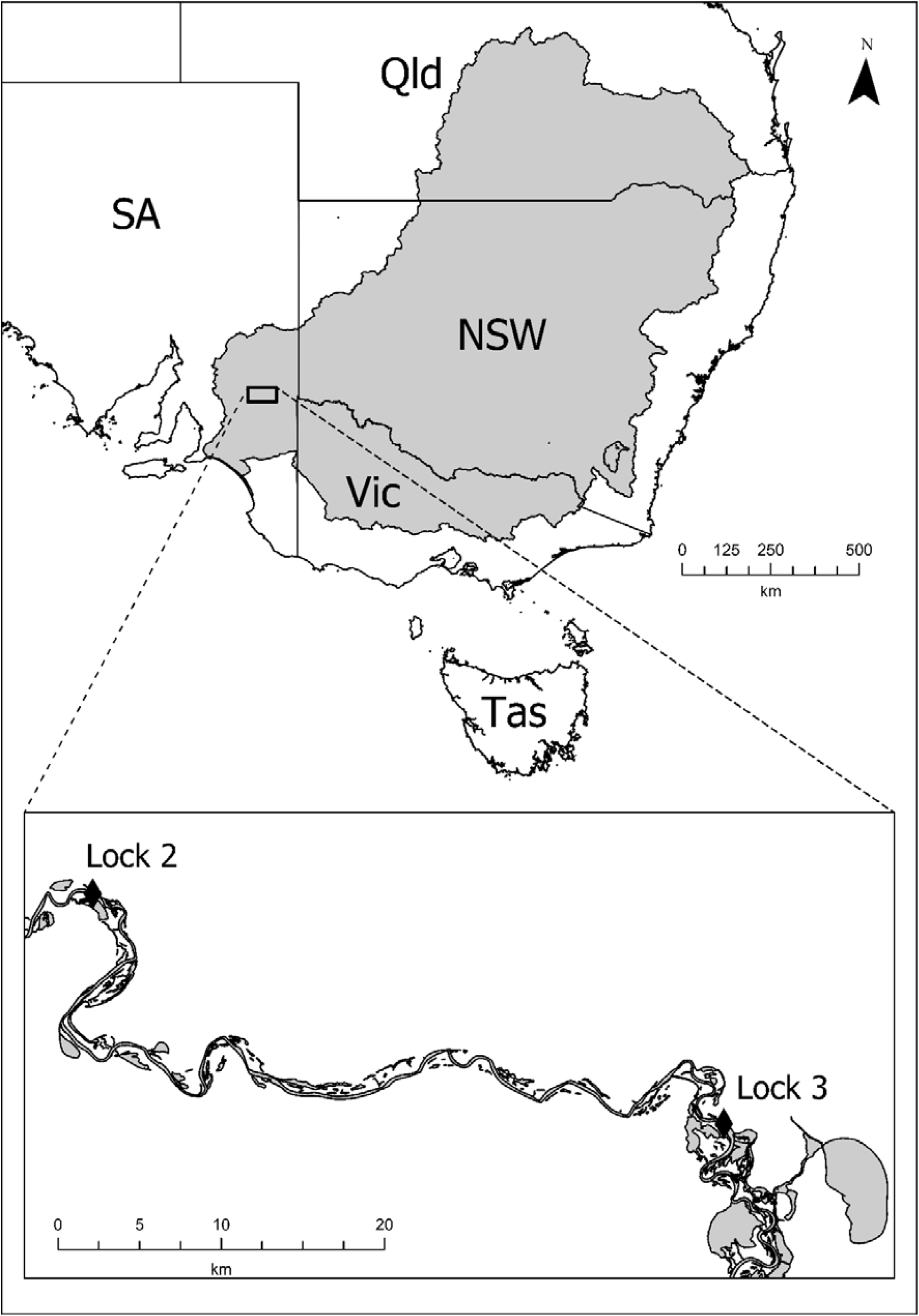
The reach between Locks 3 and 2 is at the downstream end of the Murray-Darling catchment (shaded grey). Flow is strongly influenced by regulation and extraction throughout the upstream reaches. Australian states are: SA= South Australia, Qld = Queensland, NSW = New South Wales, Vic = Victoria and Tas = Tasmania.

### Life history of southern bell frogs

The life history of *L. raniformis* is well described, with estimates for most vital rates (survival, growth, reproduction, and recruitment). Eggs hatch two to four days after laying (Anstis 2002). The likelihood of hatch (as a proxy for egg survival rate) is between 0.933 and 1.000/egg. This species ranges from north of latitude 35 °S to south of 45 °S, which spans a range of thermal conditions (from 32.9 °C maximum daily summer temperatures in the north of their range to 20.5 °C in the south). Being ectotherms, larval duration is strongly driven by temperature and as such, larval duration varies from 10–12 weeks in the north of their range to 12–15 months in the south (Anstis 2002). We used the larval duration of 70–80 days calculated at a constant water temperature of 23 °C (Cree 1984), approximately the average daily summer temperatures experienced in the study reach (which averages diurnal highs and nocturnal lows). We used the estimates of survival to metamorphosis generated for *Crinia signifera* (15–26% and 7–56%) (Williamson and Bull 1999), in the absence of species-specific estimates, these are the most-relevant estimates available.

Both sexes reach maturity in their first year (Heard et al. 2012), and females breed annually (Anstis 2017) and lay between 1885 and 4563 eggs each season (Humphries 1979, Germano and White 2008). For adult life stages we used annual adult survival probability from the closely related *Litoria aurea* (mean = 0.2172, standard deviation = 0.087) (Pickett et al. 2016). These species are similar in size, appearance, and behaviour, although *L. aurea* occurs along a more northerly latitude than *L. raniformis*. Using lines of arrested growth in the shaft of the medial phalanx to determine age (skeletochronology), *L. raniformis* can survive into its fifth year (Mann et al. 2010, G. Heard and A. Turner, Charles Sturt University, pers comm.).

### Demographic model

Within an R programming environment (R Version 4.0.2: Taking Off Again, R Project Team 2020), we constructed a stochastic population model based on a lifecycle graph (**Figure 2**) and an age-classified (Leslie) population model (Leslie 1945) that considers only females (**Figure S1**), where *f* represents fertility and *S* represents annual survival. Year 0 individuals do not breed and are not assigned a fertility. We randomly resampled fertilities (*f*) for age classes one to five years from a Normal distribution with a mean = 3245 and a standard deviation = 897 (calculated from Humphries 1979 and Germano and White 2008), which we halved to reflect the 50:50 sex ratio in this species.

**Figure 2.**
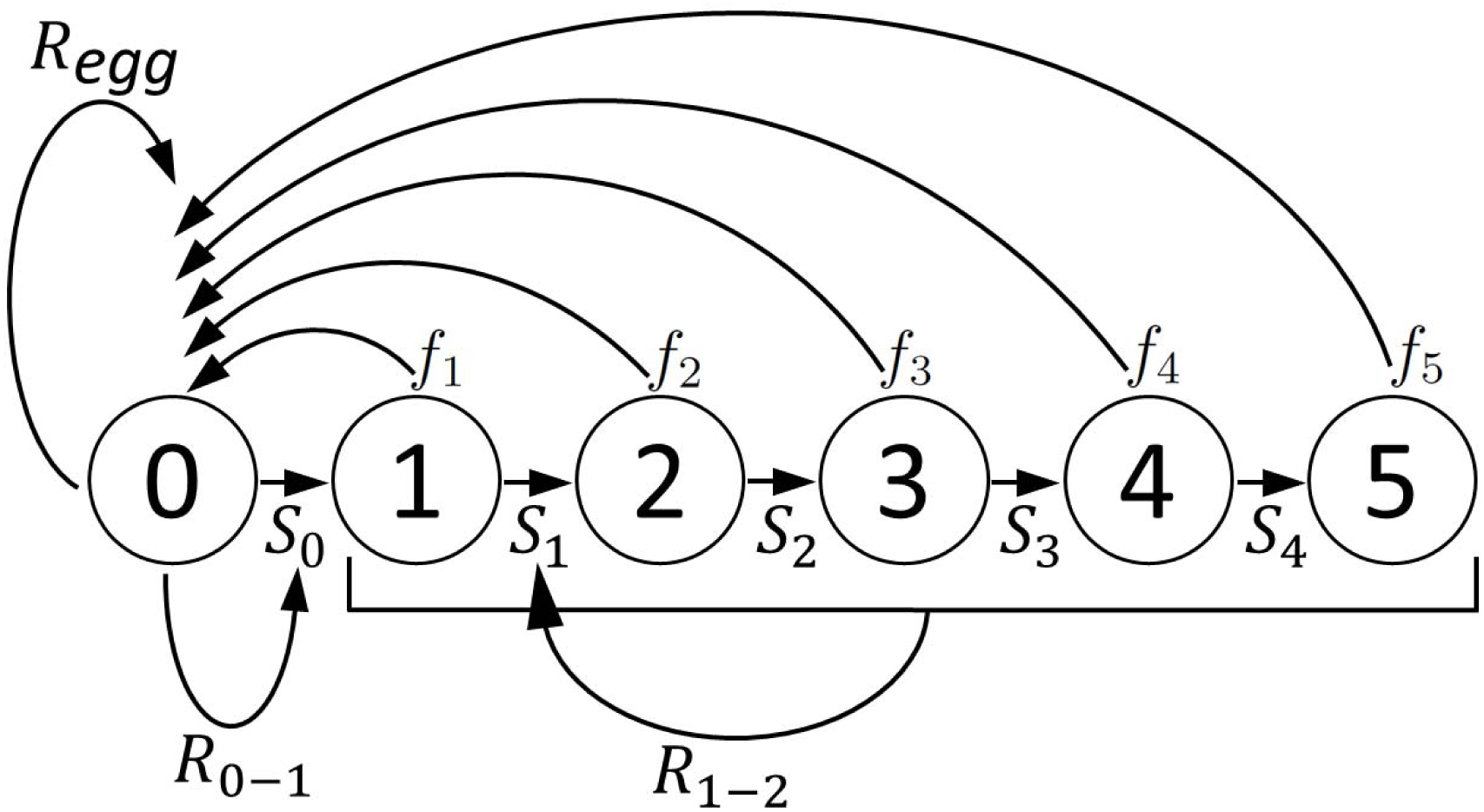
*Litoria raniformis* can live into the fifth year. Compensatory density-feedback reduction (*R*) in annual survival probability (*S*) and fertility (*f*) are calculated from both population and wetland size.

We calculated survival *S* to the end of the first year as:

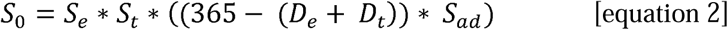

where *S*_0_ is the probability of survival from 0 to 1 year old. *S*_*e*_ is the probability of hatch (randomly resampled from a uniform distribution from 0.933–1.000). *S*_*t*_ is the probability of survival to metamorphosis, which we randomly resampled from a *β* distribution with a mean = 0.26 and a standard deviation = 0.12 (Williamson and Bull 1999). We calculated the shape parameters α and *β* of this distribution as:

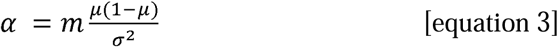

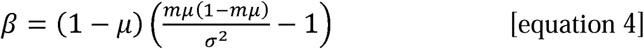

where *μ* is the mean and *σ* is the standard deviation of the *β* distribution. *D*_*e*_ is the duration of the egg stage (in days) resampled randomly from a uniform distribution between 2 and 4, and *D*_*t*_ is the duration of the tadpole stage (in days) resampled randomly from a Normal distribution between 70 and 80 days. *S*_ad_ is the daily adult survival probability calculated as:

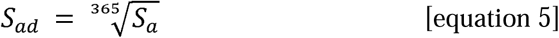

where *S*_a_ is the annual probability of survival of an adult frog sampled from a β distribution with a mean = 0.2172 and a standard deviation = 0.087 (Pickett et al. 2016). We assigned frogs in their fifth year a survival = 0, reflecting senescence and death during their fifth year (although the model permits breeding before death).

### Population sizes

We modelled *L. raniformis* populations at each of four wetland sizes (*small, medium, large, very large*). We derived these categories from typical wetlands in the region and the corresponding population size is an estimate of their respective carrying capacity using the maximum populations observed during monitoring. *Small* populations (∼ 40 individuals) reflect the carrying capacity and population dynamics present in a wetland pool several metres in diameter. A *medium* population (∼ 130 individuals) represents a wetland pool with surface area similar to 1–2 domestic swimming pools. A *large* population (∼ 300 individuals) represents a wetland pool surface area similar to an Olympic swimming pool, and a *very large* population (∼ 1000 individuals) reflects a wetland several hundred metres in diameter. The starting population of adult females was calculated as 50% of these values (as *L. raniformis* has a 50:50 sex ratio), being 20, 65, 150 and 500 female frogs, respectively. We assigned age classes for the starting female population by randomly resampling five adult survival values and then dividing the total number of females among the five age classes in these proportions. This created a more homogenous initial age structure than we might expect in a wild population. We managed this by ignoring the first 10 generations of each run as a ‘burn-in’ period, which allowed the model to stabilise to a stochastic expression of the stable-age distribution before analysis.

We allocated the initial number of eggs using an *a priori* number of spawning masses for each wetland size (*n* = 10, 30, 85, 150, respectively). We then stochastically resampled the number of eggs in each spawning mass (halved because the model only considers females). The *a priori* assignment of spawning masses did not impact population dynamics after burn-in.

### Compensatory density feedback

To stabilise long-term population growth, we incorporated three compensatory density-feedback relationships. These relationships are not available for *L. raniformis* so we estimated these functions following the form demonstrated in other anurans. First, we corrected rates of egg laying to reflect the maximum carrying capacity (*K*_egg_) at each wetland. As the number of eggs approaches the wetland’s carrying capacity, eggs are reduced following an exponential decay function of the form:

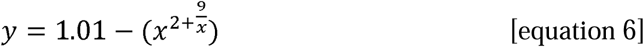

and the rate of egg reduction follows the equation:

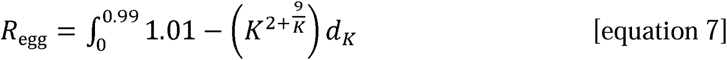

where *R*_egg_ is the total reduction in eggs laid at the wetland, *K* is the carrying capacity and *d*_*K*_ is the differential *K* (carrying capacity) (**Figure S2**). Carrying capacity is calculated for each wetland size category using the initial number of *a priori* spawning masses laid (10, 30, 85 or 150) at the maximum fecundity for the species (4563 eggs), which we halved to consider only females. In this way, frogs lay without inhibition until the total egg count approaches the carrying capacity. The highest inhibition rate corresponds to 0.99 of the wetland’s carrying capacity and all subsequent egg-laying events are reduced at this value (corresponding to a reduction of 0.0078).

The second and third compensatory density-feedback functions reduce survival during terrestrial life stages with increasing density (e.g., Harper and Semlitsch 2007, Berven, 2009). The second feedback used the total number of tadpoles present to reduce survival probabilities during the first year of life. This reflects predation and competition during larval life stages, and in the first few months post-metamorphosis. We calculated density feedback on survival probability using the function:

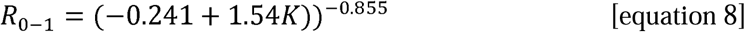

where *R*_0−1_ is the reduction factor in survival for age class 0 individuals (**Figure S3**). Here, we assigned carrying capacity *a priori* based on the size of the wetland, *small* = 200 females, *medium* = 600 females, *large* = 1800 females and *very large* = 3000 females. Density feedback on survival from 0-to 1-year age classes is applied when the total adult population > 0.3 of the wetland’s carrying capacity. We applied the highest reduction in survival at 2.1 times carrying capacity that reduces survival probability to 0.0101 (Figure S3).

The third compensatory density-feedback function applies the abundance of all adult frogs to reduce survival from the 1-to 2-year age class (*R*_1-2_) to reflect competition for resources. The form of the relationship followed the method applied for (*R*_0-1_) presented above (equation 8, **Figure S3**).

### Modelling hydrology

We considered two river-flow scenarios. The ‘regulated flow’ scenario is informed by the historical flow record immediately downstream of Lock 3 (A4260517; waterconnect.sa.gov.au). We used data starting in 1926, the year following the completion of Lock 3. River levels above the operative range of this gauge ‘drown out’ the gauge, making measurements inaccurate. During these periods, we used data from station A4260528 (6.5 km downstream of Lock 3), which is unaffected by elevated river levels. This created a continuous daily record of flow rate between Locks 3 and 2 for 83 years. The second river-flow scenario is modelled ‘natural flow’, which covers the same time period, but in the absence of extraction or regulation in the catchment (see Murray-Darling Basin Authority-MDBA 2012).

A second-order polynomial derives river heights from mean daily flow rate (Ml day^-1^) for the two flow scenarios. Based on mean daily river height (metres with respect to Australian River Height Datum, mAHD) at site A4260517 and the flow records above, the following rating-curve equation:

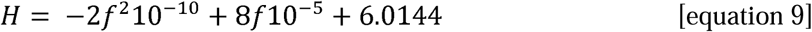

estimates river height, where *H* is river height (mAHD) and *f* is river flow (Ml day^-1^). Equation 9 represents an empirical ‘inverse rating curve’ for the site, where a rating curve is a commonly used approach to derive streamflow based on a recorded river height. We used this to create a continuous daily river height for this reach over 83 years under natural and regulated flow.

Our model exposes wetland populations to one of two states each year. A ‘wet’ year is when the wetland received sufficient water to support frog reproduction and recruitment; conversely, during a ‘dry’ year, the wetland did not receive sufficient inputs to support frog recruitment (often filling and then drying prematurely — see definition below). There is a paucity of accurate sill-height (the river height at which a wetland begins to fill) data in the reach, so rather than modelling the inundation of specific wetlands, we calculated inundation of nine possible sill heights (7, 7.5, 8, 8.5, 9, 9.5, 10, 10.5 and 11 mAHD). Following discussion with local wetland managers, we used the rule that if mean daily river height is ≥ 10 cm above sill height for ≥ 10 days during winter and spring, then the wetland is ‘wet’ and can support frog recruitment that year (K. Mason – Department for Environment and Water, Adelaide, pers comm.). It is not necessary that these ten days be consecutive to fulfil this criterion. By using this criterion, we determined which of the nine sill heights were wet, and which were dry for each of the 83 years under the natural and regulated scenarios.

### Modelling flow scenarios

We modelled 18 hydrological scenarios — these being sill heights of 7, 7.5, 8, 8.5, 9, 9.5, 10, 10.5 and 11 mAHD — each modelled under both natural- and regulated-flow conditions. To create stochastic expressions of these scenarios, we used the 83 years of daily river-height data for the flow scenario (natural or regulated flow) and classified each of those 83 years as either filling the nominated sill height (≥ 10 non-consecutive days exceeding the sill height by ≥ 10 cm during the previous winter or spring) to create a wet year, or failing to fill the sill height sufficiently and creating a dry year (when recruitment is unsuccessful). Using this sequence of 83 wet and dry years (for each specific sill height and flow combination) we created a discrete-time Markov chain (e.g., Supporting Information 1) to resample unique, stochastic, 85-year wet/dry sequences for each model scenario run.

Finally, we modelled each of the four wetland sizes (*small, medium, large, very large*) at each of nine sill heights (7, 7.5, 8, 8.5, 9, 9.5, 10, 10.5 and 11 mAHD) and each of two flow scenarios (natural and regulated flow). We ran each of these 72 wetland models stochastically 10000 times, each for 85 consecutive generations or until the population went extinct (creating a combined total of 61.2 million stochastic generations). This was sufficient to account for wetland variability in the study reach. We disregarded the first ten burn-in generations of each model and used generations 11 to 85 (or to extinction) in our analyses.

For each model, we recorded the sequence of wet and dry years from generations 11 to 85 (or extinction). For each occurrence of two or more consecutive dry years, we recorded the number of consecutive dry years and whether it resulted in local extinction. We organised these data by the average frequency of dry years at the wetland and calculated the probability of extinction for two to five consecutive dry years, noting that six consecutive dry years exceeds the maximum reproductive lifespan of the species.

To quantify the relative influence of each variable on the likelihood of local extinction we did a global sensitivity analysis on the model (Prowse *et al*, 2016). For this we modelled a large wetland with a 7.5 m sill height under regulated flow conditions and used Latin hypercube resampling to rerun 10000 discrete model iterations within the nine-dimensional parameter space (using the lhs library in R). The ranges of uncertainty used were: clutch size (1000 – 6000), duration of egg phase (0.5 – 14 days), hatch probability (0.5 – 0.98), duration of tadpole phase (50 – 90 days), mean tadpole survival (0.03 – 0.6), mean annual adult survival (0.03 – 0.35), maximum age (5 – 10 years), tadpole carrying capacity (500 – 5000 females) and juvenile carrying capacity (200 – 5000 females). We then used a boosted regression tree emulator (implemented using the dismo library in R), setting the error distribution family as Bernoulli, the bag fraction to 0.75, learning rate to 0.001, tolerance to 0.0001 and tree complexity to 2.

To examine the effect of maximum lifespan on the predictions, we also implemented a univariate sensitivity analysis on a single, large wetland with a sill height of 7.5 m under regulated flow conditions. We reran this model with senescence constrained to ages two, three, four, and five years (Appendix S1).

## Results

### ‘Wetness’ of wetlands under natural- and regulated-flow conditions

River regulation reduced the frequency of wet years by 19–34% compared to the natural-flow scenario for the same 83-year period (**Figure 3**). This equates to a reduction of wet years by 19.5% at 7 mAHD, up to 60% at 11 mAHD. This changed both the proportions and pattern of wet and dry years. Regulation increased the mean duration of successive dry years at all sill heights (**Figure 4**). Under natural conditions the mean duration of dry years did not exceed two years at any of the sill heights examined, whereas the regulated-flow scenario resulted in mean duration of dry events approaching the maximum lifespan of the species at sill heights ≥ 10 mAHD. Under natural conditions the maximum duration of dry events did not exceed the maximum reproductive lifespan of the species, except at a sill heights > 10 mAHD. The maximum duration of dry events observed under river regulation exceeded the maximum reproductive lifespan of this species at every sill height examined.

**Figure 3.**
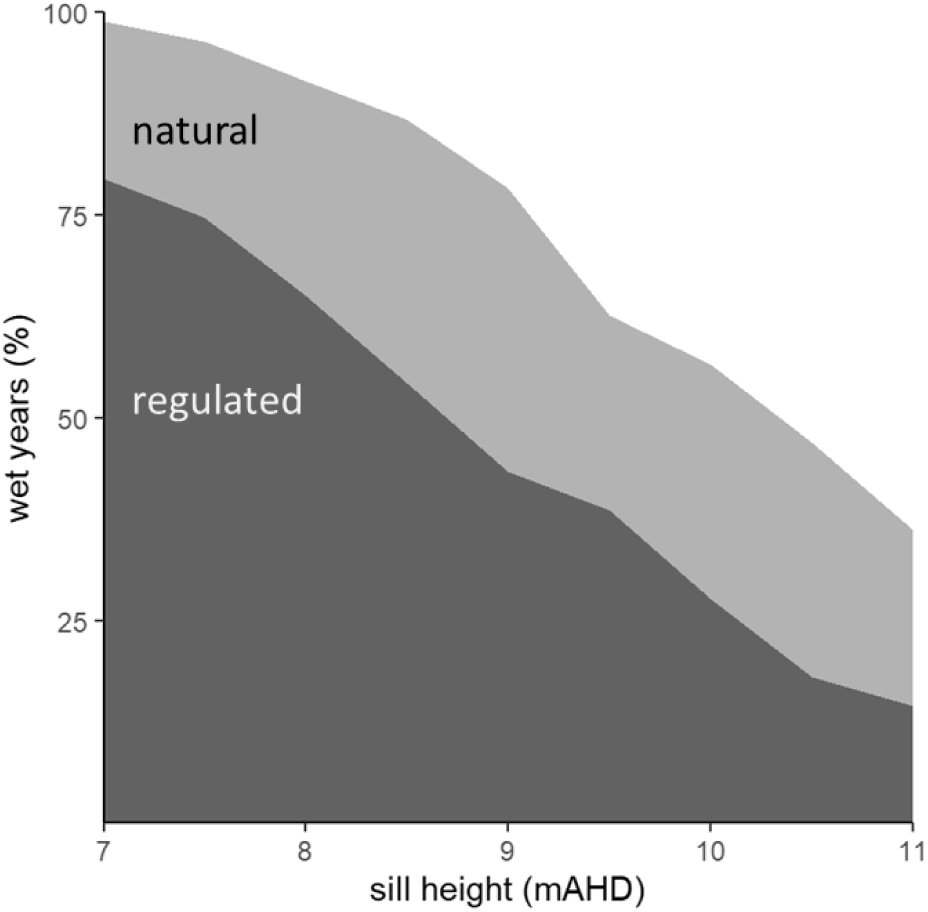
Proportion of the 83 years of observed river height (regulated) and modelled natural river height that potentially supported *Litoria raniformis* breeding (wet years) at sill heights from 7 to 11 mAHD.

**Figure 4.**
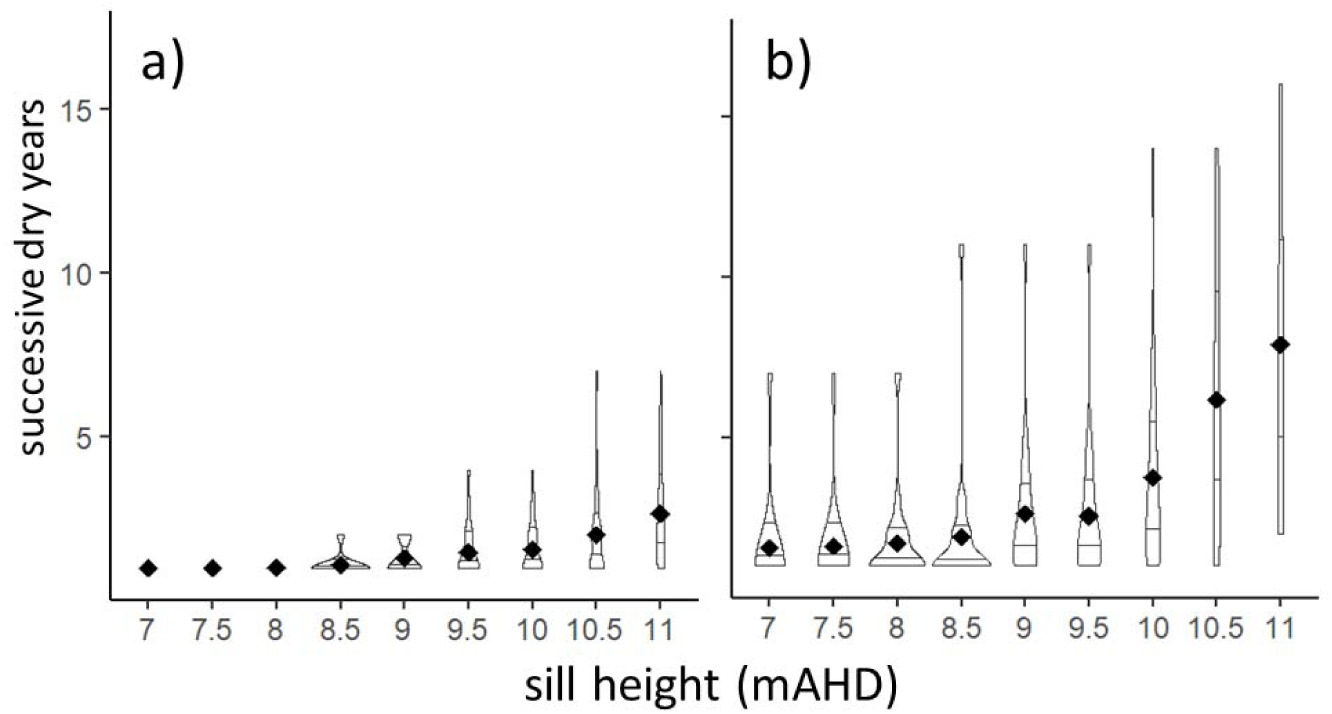
Violin plot of wetland inundation pattern over 83 years of observed data under modelled natural flow conditions (a) and river regulation (b). Sill heights are presented in metres with respect to Australian River Height Datum (mAHD). The maximum duration of dry years exceeds the maximum reproductive lifespan of *Litoria raniformis* (six years) under all regulated sill height wetlands.

### Extinction probability of each wetland scenario

Under the natural-flow scenario, the probability of extinction — Pr(Ext) — during the 85 modelled years was ∼ 0 at sill heights ≤ 8.5 mAHD (**Figure 5**). Conversely, all wetland sizes reached Pr(Ext) = 1 at sill heights ≥ 9 mAHD when flows were regulated. Thus, wetlands that historically supported *L. raniformis* populations under natural-flow conditions (i.e., ≤ 8.5 mAHD) are unreliable under river regulation. Without intervention, wetlands with sill height of ≥ 9 mAHD will not sustain *L. raniformis* populations under existing flow conditions.

**Figure 5.**
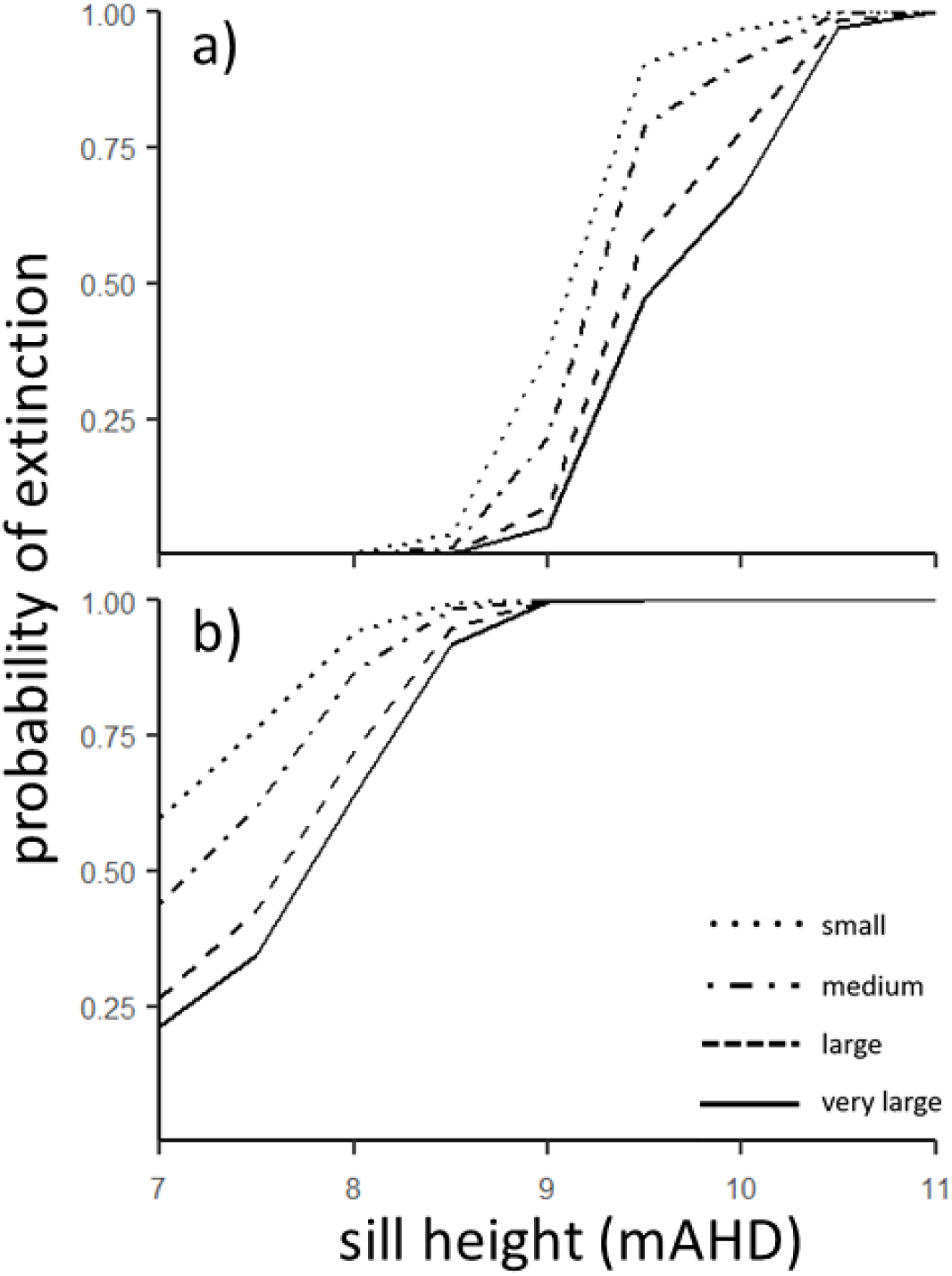
The probability of extinction at four wetland sizes under a) natural-flow and b) regulated-flow conditions. River regulation has increased the extinction probability of *Litoria raniformis* populations through most of their former habitats. These effects are strongest at smaller wetlands.

The probabilities of extinction in *very large* wetlands were 0.25 to 0.70 lower than those in *small* wetlands (**Figure 6**). This effect was more pronounced in the regulated-flow scenario than under natural-flow conditions.

**Figure 6.**
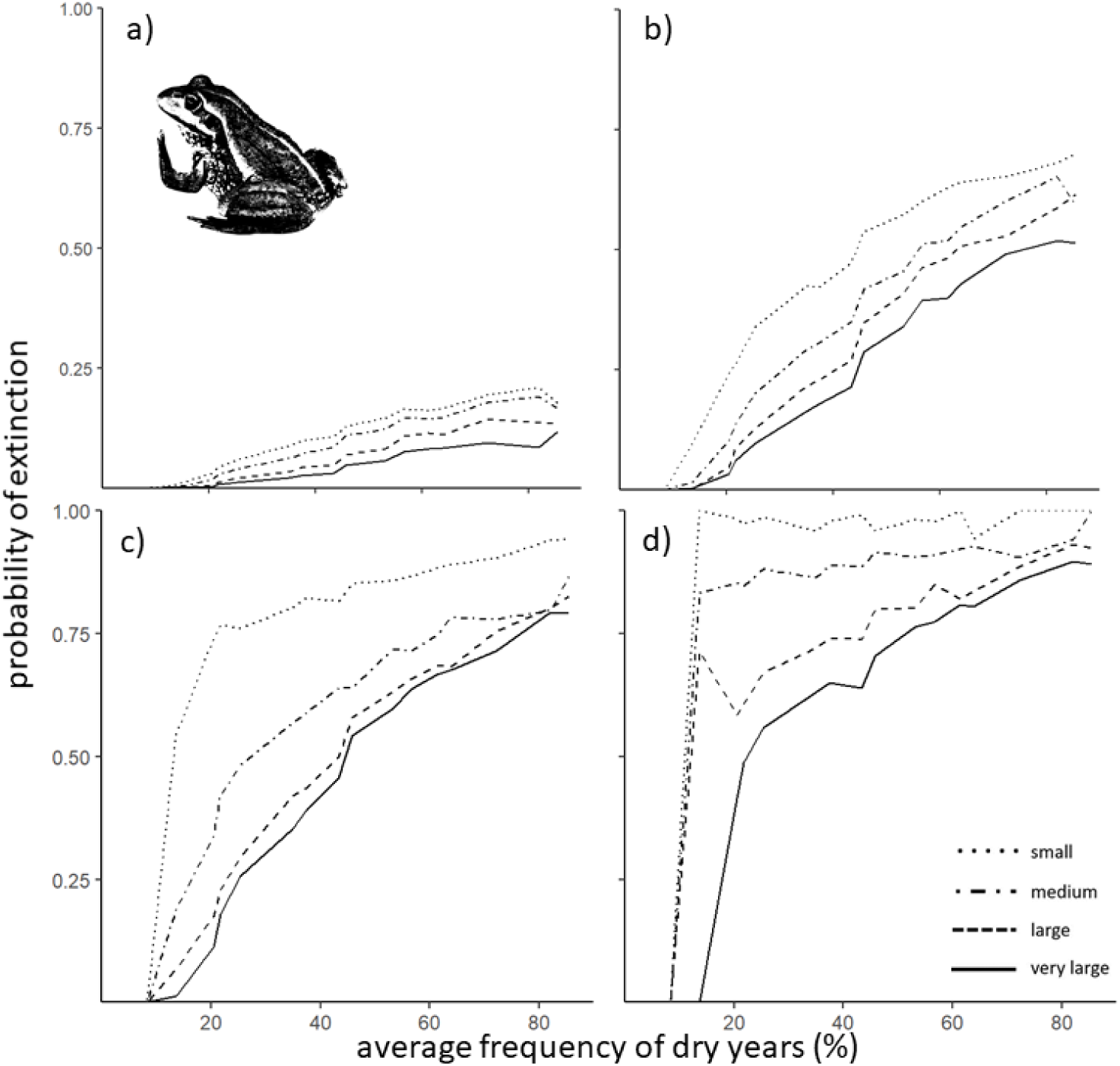
Successive dry years increase the probability of extinction and this effect increases with: increasing dry duration, decreasing wetland size, and increased average frequency of dry years at the wetland. Plots are a) 2 successive dry years, b) 3 successive dry years, c) 4 successive dry years and d) 5 successive dry years.

Two consecutive dry years resulted in Pr(Ext) < 0.25 in all treatments, including wetlands with an 80% average frequency of dry years (**Figure 6**). Increasing the number of consecutive dry years increases Pr(Ext) up to five consecutive dry years, which gives Pr(Ext) > 0.5 in all treatments except for *very large* populations with an average frequency of dry years < 20%. Smaller wetlands had increased extinction probability in all instances and the disparity between wetland sizes became more pronounced with increasing successive dry years. The average frequency of dry years at the wetland strongly influences the capacity to survive extended dry periods. Wetter sites have lower extinction probability than drier sites for each drought duration.

Boosted regression tree emulation of the nine-dimensional Latin hypercube space accounted for 96.7% of the observed variation in the model. Only two parameters strongly influenced the probability of local extinctions. These were annual mean adult survival probability, which accounted for 58.3% of the model influence and mean tadpole survival, which accounted for 21.6% of the observed variation in the model outputs (Figure 7).

**Figure 7.**
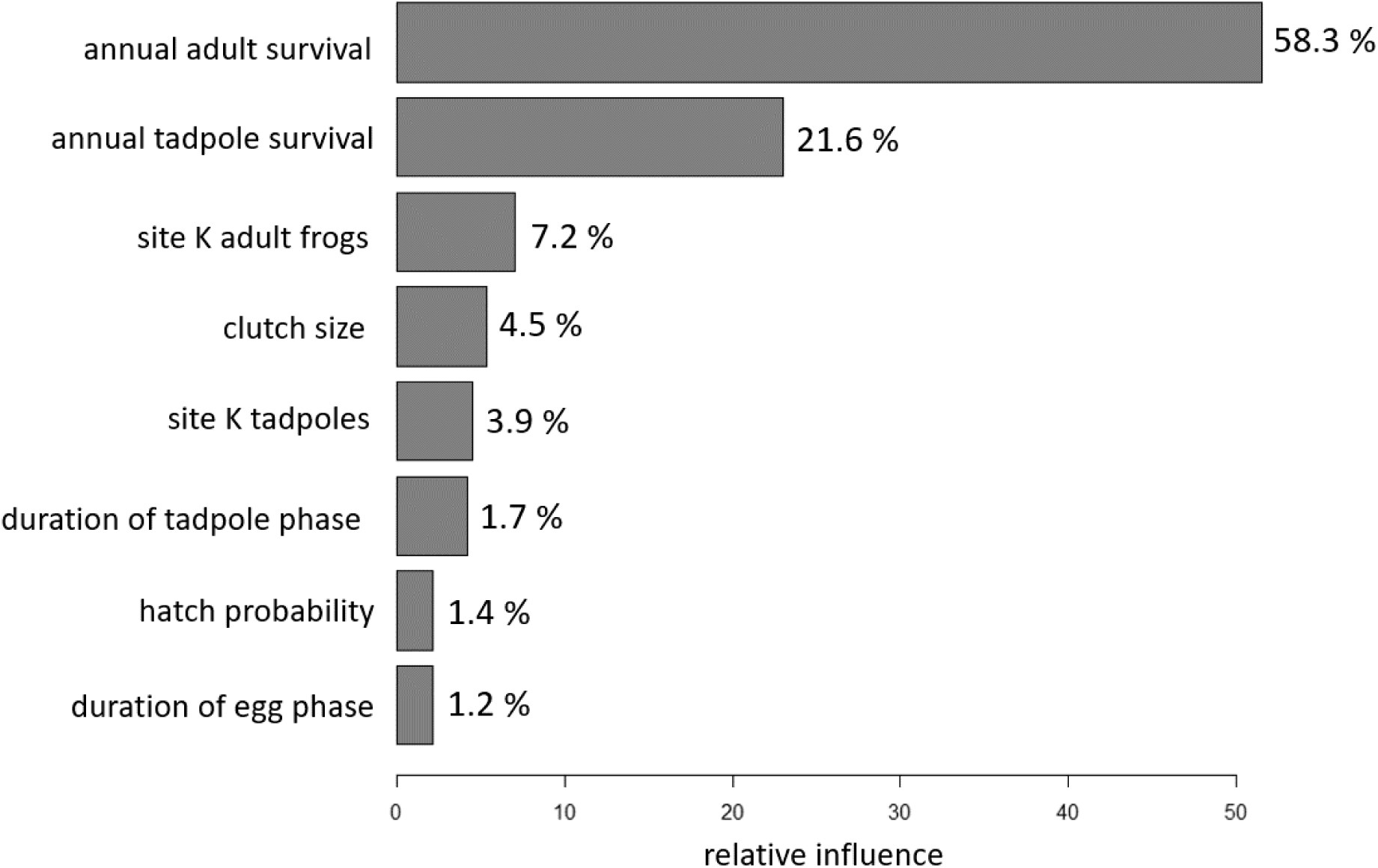
The boosted regression tree shows the relative influence of eight model parameters. Annual probability of survival in the adult and tadpole phases are the two parameters which appreciably drive model outputs.

## Discussion

### River regulation

Regulation and abstraction of flow in this catchment has resulted in wetlands that fill less often (**Figure 3**), and the average duration of successive dry years has increased by up to three years compared to natural flow conditions (**Figure 4**) (Maheshwari et al. 1995; Bice et al. 2017). For species that rely on floodplain inundation to complete their lifecycle, dry years result in reproductive failure. When breeding occurs and the site dries before completion of larval life stages, this creates population sinks that are not uncommon in amphibians breeding in ephemeral waterbodies. In isolation, reproductive failure causes a population fluctuation, but does not always appreciably increase the probability of local extinction (Taylor et al. 2006), albeit with simplification of age structure and attrition of adult populations. However, successive failures increase extinction risk, especially in short-lived species (Semlitsch et al. 1996). Without the capacity to extend lifespan through unfavourable periods (i.e., aestivation), droughts equalling or exceeding the reproductive lifespan of a species result in local extinction, a process that is likely to have driven the local extinctions of 42% of breeding sites of *Pseudophryne pengilleyi* in Kosciuszko and Brindabella National Parks in New South Wales, Australia (Scheele et al. 2012).

### *Drought and the decline of* Litoria raniformis

The flow record that informs our model includes a severe drought from 1996 to 2009. During this time, south-eastern Australia experienced a region-wide reduction in rainfall and runoff, below-average streamflow, and critical water shortages (Heberger 2011; van Dijk et al. 2013). This event, coupled with ongoing water extraction, resulted in seven or more successive dry years at all wetlands in this reach (**Figure 4**). Modelled natural flow for the same period indicates that the maximum number of successive dry years would not have exceeded four years in wetlands up to 10 mAHD (**Figure 4**). Under natural flow conditions, all wetlands < 10 mAHD could have supported *L. raniformis* through this perturbation. The persistence of some few populations through the drought is, in part, due to targeted environmental water provision during this time.

Anthropogenic shifts in the atmosphere have increased the frequency, intensity, and duration of droughts across most of the globe and these trends are likely to continue (Chiang et al. 2021). Droughts exacerbate the ecological impacts of river regulation and scientifically defensible approaches to the manipulation of aquatic resources (like those presented here) will become increasingly necessary to conserve some freshwater species, particularly wetland- and pond-breeding amphibians.

### Model assumptions

Our population model relies on two main assumptions: that (*i*) *L. raniformis* breed in their fifth year before senescence, and (*ii*) there is no population exchange between wetlands. However, our boosted regression tree analysis (**Figure 7**) and single-parameter perturbation (sensitivity) analysis of age (**Appendix S1**) show little influence of maximum age on extinction probability. The strong influence of survival probabilities (**Figure 7**) results in a demography that is weighted strongly towards younger age classes and this first assumption is unlikely to skew the results if incorrect. Similarly, the model outputs are unlikely to be affected by inaccurate calculation of carrying capacities, clutch size, hatch probability, or duration of egg or tadpole life stages (**Figure 7**).

The second major assumption is that wetlands are isolated and lack immigration or emigration. As such, modelled local extinctions are an endpoint after which recolonisation cannot occur. Amphibians are physiologically dependent on moist environments, have relatively poor dispersal capacity (compared to other tetrapods), and can show strong site fidelity, and these traits suggest limited capacity for recolonisation (Blaustein et al. 1994). Despite limited dispersal, amphibians are frequently thought to exist in metapopulations (Levins 1969; Smith and Green 2005). We propose that *L. raniformis* in this system meets the four characteristics of a metapopulation (Hanski et al. 1995): (*i*) habitat patches support local breeding populations, (*ii*) no single population is large enough to ensure long-term survival, (*iii*) patches are not too isolated to prevent recolonisation, and (*iv*) local dynamics are sufficiently asynchronous to make simultaneous extinction of all local populations unlikely. Despite the semi-arid/arid climate, *L. raniformis* can move hundreds of metres in a single night and can colonise over distances of 500 m (Herbert 2000; Department for Environment and Heritage, 2004; Wassens et al. 2008). We anticipate that dispersal between favourable patches (albeit infrequent and increasingly difficult in this system due to fragmentation, river regulation, and aridification) can result in recolonisation and contributes to the persistence of this species at the landscape level through the rescue effect (Brown and Kodric-Brown, 1977). We are developing metapopulation models for this reach to further explore these dynamics.

Our methodology takes a cautionary approach to conservation. We deliberately model wetlands without the stabilising effects of metapopulation structure. We acknowledge this approach produces higher extinction probabilities than we might expect from wetlands within a network, but this also produces cautionary conservation advice that supports wetlands without reliance on stochastic phenomena. In systems where populations have become fragmented, successful movement between populations is low, or movements vary broadly between years, we recommend this approach to prioritise the active management of threatened amphibians.

### Environmental water provision

In highly regulated catchments, restoring large-scale historic inundation patterns is neither possible nor desirable. However, providing specific reaches and wetlands with environmental water can recreate critical components of historic flow regimes. The assumption is that reconstructing specific components of ‘natural’ flow can maintain key ecosystem processes through the timely, sequential ecological cues inherent in ‘natural’ flow (Poff et al. 1997). In highly modified and non-stationary conditions, a more mechanistic understanding of environmental water requirements is valuable (Poff 2018). An alternative approach uses thresholds of inundation frequency, and in turn extinction probability to prioritise the delivery of a ‘designer’ flow regime. When well-designed, this approach generates efficiencies in the volumes of water delivered while still eliciting the desired ecological response (in this case, a reduction in the probability of extinction for target amphibian populations).

Our results show that water deliveries maintaining no more than one sequential dry year will reliably support *L. raniformis* (**Figure 6**). For wetlands with low extinction probability, or when environmental water budgets are limiting, water provision to ensure no more than two successive dry years will maintain extinction risk < 0.25 in all wetland sizes. Using this approach, a rotating roster of wetlands could be watered to offset individual wetland risk, noting that local extinctions will likely rely on dispersal from neighbouring wetlands to re-establish. The highest priorities for intervention are wetlands that have experienced four or five years without recruitment and where *L. raniformis* is still present (**Figure 6**). These populations should be watered during the next breeding season to reduce the probability of local extinction. These priorities are not intended to be cycled in perpetuity, especially in situations where extinction risk is high. For example, a wetland with a 50% extinction probability after five dry years could require several favourable years to recover before it could be expected to persist through a second five-year drought.

Our model makes three main assumptions regarding wet years that influence the techniques for environmental water delivery: that 1. wetlands support adult frogs between breeding seasons, 2. breeding occurs during each wet year, and 3. wet years (and particularly successive wet years) do not accumulate fish and invertebrate predators. Water delivery should be tailored to meet these assumptions, including maintaining small pools to allow adult frogs to rehydrate during dry periods, watering to support components of the vegetation community that are important for breeding, and periodic drying to reduce predator densities. This last point is particularly important as many systems do not dry naturally between wet events and will accumulate predators more readily than this example. When modelling these systems, an additional relationship is required which incorporates the effects of predator densities. Caution is also required when designing flows to ensure the delivery method does not damage existing habitat features, e.g., erosion or scouring of breeding habitats with high delivery velocities (Kupferberg, 1996).

## Conclusions

Our approach informs thresholds for environmental water provision that can be applied where demographic data are available and where a clear relationship exists between intervention and recruitment — for example, the delivery of water to extend hydroperiod in *Rana sevosa* (Seigel et al. 2006) or *P. pengilleyi* (Scheele et al. 2012). This approach is not specific to amphibians, nor to environmental water provision, and can also be applied where episodic events are directly linked to reproductive outcomes, such as fire intermittency and germination in pyrophytic plants, or supraseasonal flooding events in arid-zone seed germination.

Environmental water provision to amphibian breeding habitats can reduce the probability of extinction in populations with insufficient inundation frequency. We show that stochastic population models, typically constructed for population viability analyses, can be used to determine critical hydrological thresholds, and that these thresholds are sufficiently nuanced to prioritise intervention locations and guide the frequency of environmental water provision. Strategic *a priori* planning and efficient water use will become increasingly important techniques in amphibian conservation, especially in regulated river systems.

## Acknowledgements

We thank the Murraylands and Riverland Landscapes Board and the Nature Foundation SA for their generous support and the Murray-Darling Basin Authority for providing time-series data of natural flow.

## Declarations

### Ethics approval

Not Applicable

### Consent to participate

Not Applicable

### Consent for publication

All authors consent to the publication of this manuscript, its supplementary information and the associated code.

### Availability of data and material

There are no novel datasets associated with this work. All data are cited and referenced herein

### Code availability

Data and code available from: https://github.com/RupertLovesEcology/RiverRegulation_Frog_PopModel

### Authors’ contributions

RM and MSG completed the hydrological modelling, RM and CJAB designed and constructed the population models. All authors contributed to the manuscript and provided editorial advice. This article does not contain any studies with human participants or animals performed by any of the authors.

## Supporting Information

Additional Supporting Information can be found in the online version of this article.

**Appendix S1**. To examine the effects of maximum lifespan on the predicted probability of extinction, we applied a single-parameter perturbation (sensitivity) analysis on a single modelled scenario (a large wetland with a sill height of 7.5 metres with respect to the Australian Height Datum (mAHD) under a river regulation flow scenario).

**Figure S1**. The Leslie matrix (L1) assigns fertility to age classes 1 – 5 in the top row and age-specific annual survival probability on the sub-diagonal.

**Figure S2**. Inhibition of egg laying at a wetland follows an exponential decay function. If the total number of eggs laid is greater than 0.8 of the wetland’s carrying capacity, subsequent eggs are reduced at an increasing rate.

**Figure S3**. Compensatory density feedback on survival rate of eggs to the 1-year age class and survival of the 1-year age class to the 2-year age class based on an exponential decay function. Reduction of survival probability starts when the population exceeds 0.3 of carrying capacity.

